# Two extreme Loss-of-Function *GRIN2B*-mutations are detrimental to tri-heteromeric NMDAR-function, but rescued by pregnanolone-sulfate

**DOI:** 10.1101/2022.12.13.520218

**Authors:** Shai Kellner, Shai Berlin

## Abstract

Mutations within various N‐methyl-D-aspartate receptor (NMDAR) subunits are tightly associated with severe pediatric neurodevelopmental disorders and encephalopathies (here denoted *GRINopathies*), for which there are no treatments. NMDARs are tetrameric receptors and can be found at the membrane of neurons in various compositions, namely in di- or tri-heteromeric forms. The GluN2B subunit appears very early in development and, therefore, prenatally this subunit is predominantly found within di-heteromeric receptors, exclusively composed of the GluN1 and GluN2B subunits. Postnatally, however, the GluN2A subunit undergoes rapid increase in expression, giving rise to the appearance of tri-heteromers containing the GluN1, GluN2A and GluN2B-subunits. The latter are emerging as the principal receptor-type postnatally. Despite more than a decade of research of numerous *GRINopathies*, not much is known regarding the effect of *GRIN* variants when these are assembled within tri-heteromers. Here, we have systematically examined how two *de novo GRIN2B* variants (G689C and G689S) affect the function of di- and tri-heteromers. We show that whereas a single mutated subunit readily instigates a dominant negative effect over glutamate affinity of tri-heteromers, it does not dominate other features of the receptor, notably potentiation by pregnanolone-sulfate (PS). This led us to explore PS as a potential treatment for these two severe loss-of-function (LoF) mutations in cultured neurons, in which case we indeed find that the neurosteroid rescues current amplitudes. Together, we present the first report to examine LoF *GRIN2B* mutations in the context of di- and tri-heteromeric receptors. We also provide the first demonstration of the positive outcome of the use of a *GRIN2B*-relevant potentiator in the context of tri-heteromers. Our results highlight the importance of examining how different mutations affect features in various receptor subtypes, as these could not have been deduced from observations performed on purely di-heteromers. Together, our study contributes to the ongoing efforts invested towards understanding the pathophysiology of *GRINopathies* as well as provides insights towards a potential treatment.

## Introduction

N-methyl-D-aspartate receptors (NMDARs or GluNRs) are glutamate-gated receptors, commonly expressed at postsynaptic loci^1^, where they play essential roles in synaptic maturation and refinement, dendritogenesis, plasticity, learning and memory, to name a few^2–7^. Owing to their quintessential roles in the central nervous system across development, mutations within *GRIN* genes (of which there are seven: *GRIN1, GRIN2A-D, GRIN3A-B*)^2,3^ are, as of 2010^8^, intimately associated with neurodevelopmental disorders, severe mental retardations, epilepsy, and more^8–12^. Since the emergence of the first report by Endele et al.^8^, systematic screenings for NMDAR mutations in pediatric patients has unearthed thousands of inherited or *de novo GRIN* mutations^13^; the majority of which are concentrated within *GRIN2A* and *GRIN2B* (46% and 38%, respectively)^9,13–19^. Unfortunately, whether and how most mutations affect channel function and/or expression is unknown, and the rate at which new variants are emerging in the literature (mirroring the clinic) exceeds the rate at which these can be functionally characterized and screened for responsiveness to NMDAR-selective drugs. Consequently, only a fraction of mutations has been characterized, and in most cases (including our own efforts), characterization is almost exclusively restricted to one type of receptor-composition, namely di-heteromeric form (see below).

NMDARs are very large tetrameric complexes assembled from two obligatory GluN1-subunits with two other, similar or dissimilar, GluN2 and/or GluN3-subunits, denoted di- and tri-heteromeric channels, respectively^2,20^. Prenatally, the most common form in the forebrain is the di-heteromeric GluN1/GluN2B-receptor, whereas tri-heteromeric GluN1/GluN2A/GluN2B-receptors are emerging as the most predominant subtype, postnatally through adulthood, particularly in the hippocampus^21–24^. These thereby suggest that, in patients with heterozygous *GRIN* mutation, mutated variants should exist in multiple (pure or mixed) di- and tri-heteromeric channel types^25^. How, and whether, single mutated subunits can affect the function of various forms of the receptors remains poorly understood, owing to the handful of publications on this subject (e.g., ^11,24,25^).

To bridge this gap, here we specifically examine the effect of two *extreme* loss-of-function (LoF) GluN2B-mutations (p.G689C and p.G689S^9^) in the context of di- and tri-heteromeric channels. We have previously shown that both mutations reside at the entry of the ligand-binding domain (LBD) of the GluN2B subunit and, thereby, cause a dramatic reduction in glutamate affinity (up to ∼2000-fold) in receptors specifically containing two copies of the mutant GluN2B variant^9^. Under normal synaptic transmission, these channels are thereby completely silent. However, whether a single mutated subunit could affect *mixed* di-heteromeric (i.e., a receptor containing a mutated and a *wt* GluN2B subunit) or tri-heteromeric channels (i.e., containing a mutated GluN2B and a *wt* GluN2A subunit) remains unknown. To address this, we employ a unique ER-retention technique^23^ to limit expression of a desired receptor-population at the membrane of cells. We find that the inclusion of a single GluN2B-variant within *mixed* di- or tri-heteromeric channels is sufficient to prompt a strong reduction in the receptors’ glutamate affinity, but these reductions are not as drastic as in purely di-heterometric receptors containing two copies of the variants. This observation is supported by the ability of a GluN2B-selective potentiator (spermine) to potentiate *mixed* di-heteromeric channels. Interestingly, and counterintuitively, we find that the variants have no negative effect over the responses of the channels to pregnenolone-sulfate (PS), a GluN2A and 2B-selective potentiator. Indeed, application of PS rescues the amplitude of NMDA-dependent currents in neurons expressing the variants individually. Together, our results show the difficulty to predict *a priori*, whether, and how, each variant may affect features of the most abundant tri-heteromeric form of the receptors. Finally, we suggest that PS may serve as a potential treatment for LoF mutations, even in the case of extremely deleterious mutations, such as G689C and G689S.

## Results

### Loss-of-Function mutations in GluN2B instigate a dominant-negative effect over mixed di-heteromeric GluN2B-containing receptors

We have previously identified two analogous *de novo* mutations in the ligand binding domain (LBD) of GluN2B at residue G689 (G689C and G689S^9^). Both mutations instigate a dramatic reduction in glutamate affinity (EC_50_) in *pure* di-heteromeric receptors containing two mutant GluN2B-subunits (GluN2B*) co-assembled with two GluN1a-wt subunits (for brevity, we omit mention of GluN1 subunits hereafter). Interestingly, we also found that the mutations affect other receptor functions that are regulated by other parts of the subunit. However, owing to the heterozygous-nature of most *GRINopathies*, GluN2B*-subunits can also multimerize with GluN2B-*wt* subunits to form *mixed* di-heteromeric channels, which is likely a better reflection of the expression patterns *in-vivo*^7,25^. To try to examine the effect of the mutations on *mixed* di-heteromeric receptors, we sought means to control channel stoichiometry at the membrane; leading us to explore a previously reported selective cell-surface-expression method for GluNRs^23,26^. Briefly, we tagged the carboxy termini of different GluN2B-subunits with leucine zipper motifs from GABA_B1_ or GABA_B2_ along ER-retention motifs (denoted C1 and C2, respectively) (see **methods**), while leads to the exclusive surface expression of receptors composed of two different C1- and C2-tagged GluN2-subunits (denoted C1/C2), whereas C1/C1 or C2/C2-containing channels are retained in the ER (**Fig. 1a**).

**Figure 1.**
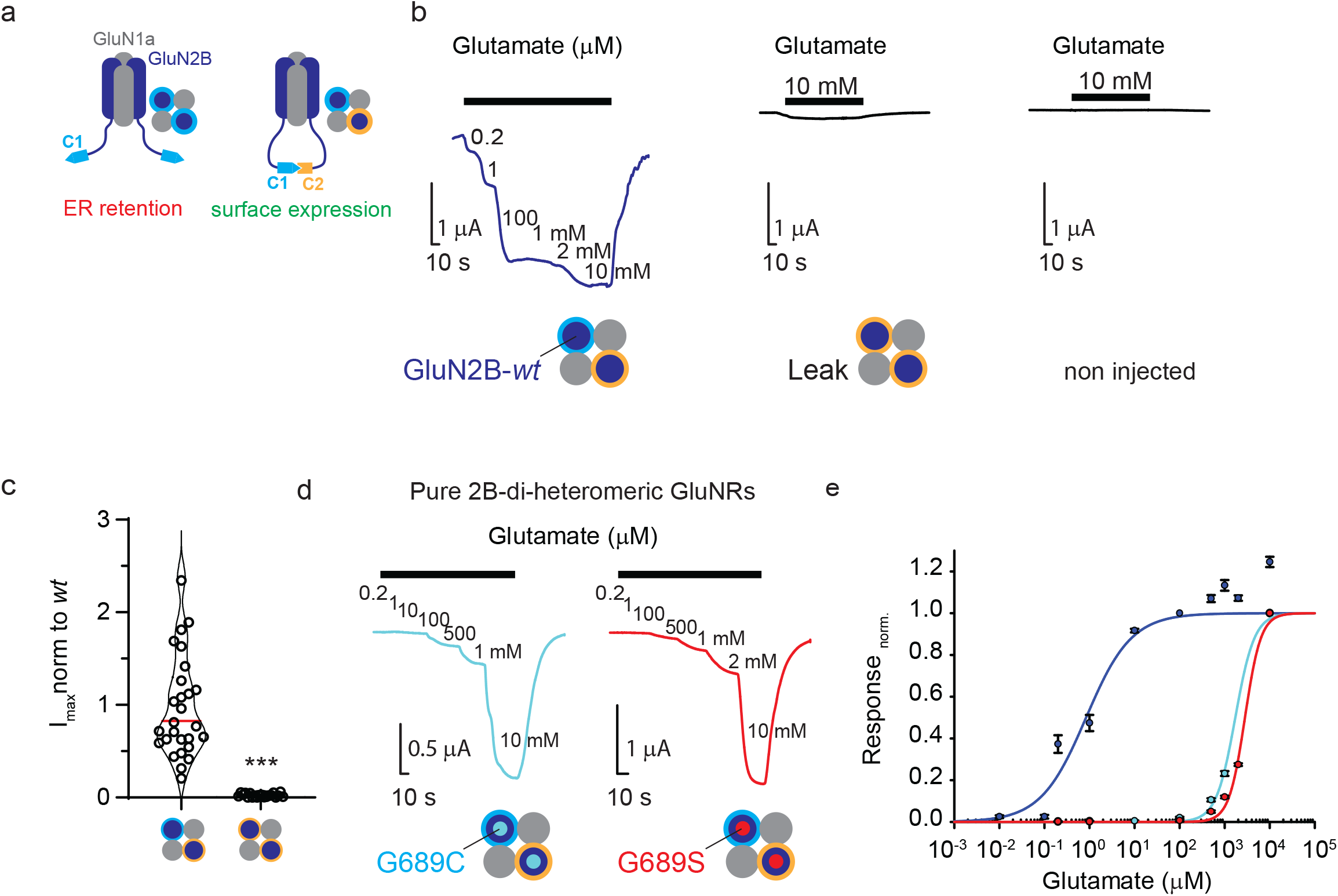
Selective expression of GluNR-subtypes at the surface of *xenopus laevis* oocytes. **a**. Cartoon depiction of a di-heteromeric receptor composed of two wildtype GluN1a subunits (gray) assembled with two GluN2B-subunits (dark blue). Left: receptors containing two GluN2B-subunits tagged with C1 tail (cyan carboxy-termini outlines dark blue circle) are retained in ER, whereas receptors containing two GluN2B subunits with C1-(cyan) and C2-termini (orange) are trafficked to the membrane (right). **b**. Representative traces showing glutamate-dependent currents recorded from oocytes co-expressing GluN1a-*wt* with GluN2B*wt*-C1/C2 (left, dark blue trace), or just with GluN2B*wt*-C2 (middle; leak); non-injected oocytes (right). Glutamate (and glycine) application is noted by black bar above traces. **c**. C1/C2-containing receptors yield large currents easily discernable from leak currents. Summary of four independent experiments showing maximal currents (I_max_) comparing GluN2B-wt-C1/C2 (n= 34) and leak currents (from oocytes expressing GluN2B*wt*-C2, n= 20). Median is highlighted in red. **d**. Representative traces showing glutamate dose-response currents recorded from oocytes co-expressing GluN1a-*wt* with GluN2B-G689C-C1/C2 (cyan trace; G689C mutation is denoted by a small filled cyan circle within the GluN2B subunit-dark blue) or with GluN2B-G689S-C1/C2 (red trace; G689S-red filled circles). **e**. Summary of dose-response curves for GluN1a-*wt*+GluN2B*wt*-C1/C2 (blue), GluN1a-*wt*+GluN2B-G689C-C1/C2 (cyan) and GluN1a-*wt*+GluN2B-G689S-C1/C2 (red).

We thereby produced GluN2B-G689C and GluN2B-G689S subunits tagged with C1 or C2, co-expressed these along GluN1a-*wt* subunit in *Xenopus* oocytes, and assessed the activity of different GluNR compositions via two-electrode voltage clamp (TEVC) (see **methods**)^27,28^. We initially co-expressed GluN2B-*wt*-C1 with GluN2B-*wt*-C2 and find that these channels readily express at the membrane and yield specific glutamate-dependent currents (**Fig. 1b, c**), whereas the sole expression of just one subunit (e.g., GluN2B-*wt*-C2 with GluN1a) yields very small, albeit *bona-fide*, glutamate-currents (denoted leak), but these were mainly noticeable when oocytes are exposed cells to high (saturating) glutamate concentrations (10 mM, **Fig. 1b, c, but also see Suppl. 1 for more leak examples**). We next proceeded to compare the apparent glutamate affinity (EC_50_) of purely di-heteromeric channels composed of GluN2B-G689C-C1/C2 and GluN2B-G689S-C1/C2-channels, to GluN2B-*wt-*C1/C2 receptors. We observed that GluN2B-G689C and GluN2B-G689S instigate a drastic reduction in EC_50_, namely ∼2000- and >3000-fold reduction in EC_50_, respectively, on par with our previous observations with the non-tagged variants (**Fig. 1d, e Table 1**)^9^.

**Table 1.** Spermine used at 200 μM, PS-100 μM. Significance was assessed by comparing to control group in each section (by shades). *, p < 0.05; ***, p<0.001; n.s., not significant

| Group | Glutamate<br>EC<sub>50</sub><br>(taken from<sup>9</sup>) | Glutamate<br>EC<sub>50</sub> | Spermine<br>Potentiation<br>(%) | PS<br>Potentiation<br>(%) |
| --- | --- | --- | --- | --- |
| Purely di-heteromeric<br>GluN2Bwt | 1.4±0.04 µM<br>(N=3, n=43) | 0.85±0.4 µM<br>(N=6, n=38) | 97±5.6<br>(N=3, n=25) | 203±16%<br>(N=3, n=16) |
| Purely di-heteromeric<br>GluN2B-G689C | 1.54±0.14 mM<br>***<br>(N=2, n=31) | 1.72±0.16 mM<br>***<br>(N=2, n=13) | - | 296±14%<br>(N=3, n=18)<br>*** |
| Purely di-heteromeric<br>GluN2B-G689S | 2.56 ± 0.40 mM<br>***<br>(N=2, n=23) | 2.82±0.24 mM<br>***<br>(N=2, n=9) | - | 368±18%<br>(N=3, n=23) *** |
| Mixed di-heteromeric<br>GluN2Bwt/GluN2B-G689C |  | 368.1±55.6 µM<br>***<br>(N=6, n=54) | 78.5±5.15<br>(N=2, n=20)<br>*** | 177±15%<br>(N=3, n=13)<br>n.s. |
| Mixed di-heteromeric<br>GluN2Bwt/GluN2B-G689S |  | 760±64µM<br>***<br>(N=1, n=13) | 61±3.31<br>(N=2, n=24)<br>*** | 199±10%<br>(N=3, n=20)<br>n.s |
| Tri-heteromeric<br>GluN2Awt/GluN2Bwt |  | 1.54±0.64 µM<br>(N=3, n=33) | -26±1.9<br>(N=2, n=12) | 89±4.3%<br>(N=4, n=28) |
| Tri-heteromeric<br>GluN2Awt/GluN2B-G689C | | 502±88 μM<br>***<br>(N=1, n=10) | -30.1±2.7%<br>(N=1, n=10) | 112±9.6%<br>(N=3, n=13) n.s |
| Tri -heteromeric<br>GluN2Awt/GluN2B-G689S | | 936±91 μM<br>***<br>(N=1, n=11) | -25±2.33%<br>(N=2, n=19) | 101±6%<br>(N=2, n=13) n.s |

Spermine used at 200 μM, PS- 100 μM. Significance was assessed by comparing to control group in each section (by shades). \*, $p < 0.05$ ; \*\*\*, $p < 0.001$ ; n.s., not significant

We then proceeded to examine the effect of single variants within a *mixed* di-heteromeric channel (i.e., GluN2B-*wt*-C1+GluN2B*-C2). Expectedly, we noted that *mixed* channels showed major reductions in EC_50_ values, with GluN2B-G689S-containing channels displaying a more severe rightward shift (**Fig. 2a, b, Table 1**). Nevertheless, both mixed di-heteromeric channels showed ∼four-fold higher (i.e., improved) affinities compared to purely di-heteromeric mutated channels (**Fig. 2b, solid vs. dashed plots, and see Table 1**). These observations are somewhat reminiscent of our previous report, in which we have also observed slightly improved EC_50_ values in oocytes injected with different ratios of *wt* and mutated subunits’ mRNAs (see ^9^).

**Figure 2.**
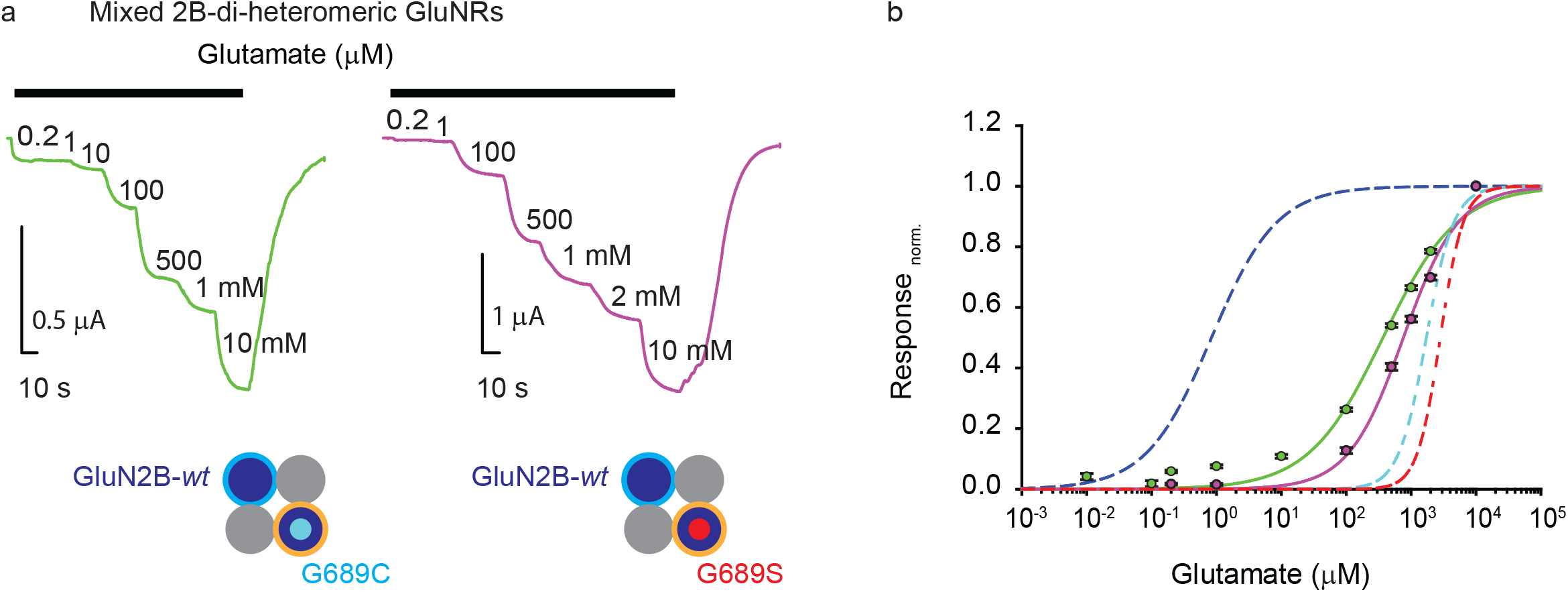
GluN2B-variants reduce glutamate potency of *mixed* di-heteromeric receptors. **a**. Representative traces from oocytes expressing *mixed* di–hetereomers composed of GluN1a-*wt* with GluN2B*wt*-C1 and GluN2B-G689C-C2 (lime trace) or with GluN2B-G689S-C2 (purple trace), in response to increasing glutamate concentrations. Glutamate (and glycine) application is noted by black bar above traces. Glutamate concentrations (in μM) are noted next to current steps. mM units are explicitly noted next to steps. **b**. Summary of dose-response curves for *mixed* di–heteromeric GluN2B-receptors, colored-coded as in **(a)**. Dashed lines depict dose-response curves for purely di-heteromeric receptors shown in **Fig. 1e**: GluN2B*wt*-C1/C2 (blue), GluN2B-G689C-C1/C2 (cyan) and GluN2B-wt-C1/C2 (red).

Thus, our current results demonstrate that a single variant within a mixed di-heteromeric channel is sufficient to engender a strong dominant negative effect over the entire channel, however the variant’s affinity does not govern the apparent affinity of the entire complex (i.e., is not the limiting factor). Instead, the channel’s EC_50_ is adjusted by both subunit types (*wt* and mutant). Our results thereby elegantly demonstrate that robust channel opening (on a macro-current level) depends on the *liganding* of all subunits, strongly supporting previous observations^29–31^, as well as demonstrates the positive allosteric effects between subunits^32^.

We next examined the effect of the variants over the more common, if not the dominant, channel-subtype in the adult brain, namely tri-heteromeric channels composed of GluN2A and GluN2B-subunits^7^. GluN2A-*wt* tagged with the C1 tail, co-expressed with C2-tagged GluN2B variants, also exhibited severe LoF; yielding ∼300 and ∼600-fold reductions in EC_50_ by GluN2B-G689C and 2B-G689S variants, respectively, in comparison to GluN2A/2B-*wt* tri-heteromeric channels (**Fig. 3, Suppl 1a-b, and summary in Table 1**). Together, our results demonstrate that GluN2B-variants induce severe reductions in glutamate potency when incorporated in either *mixed* di- or tri-heteromeric channels.

**Figure 3.**
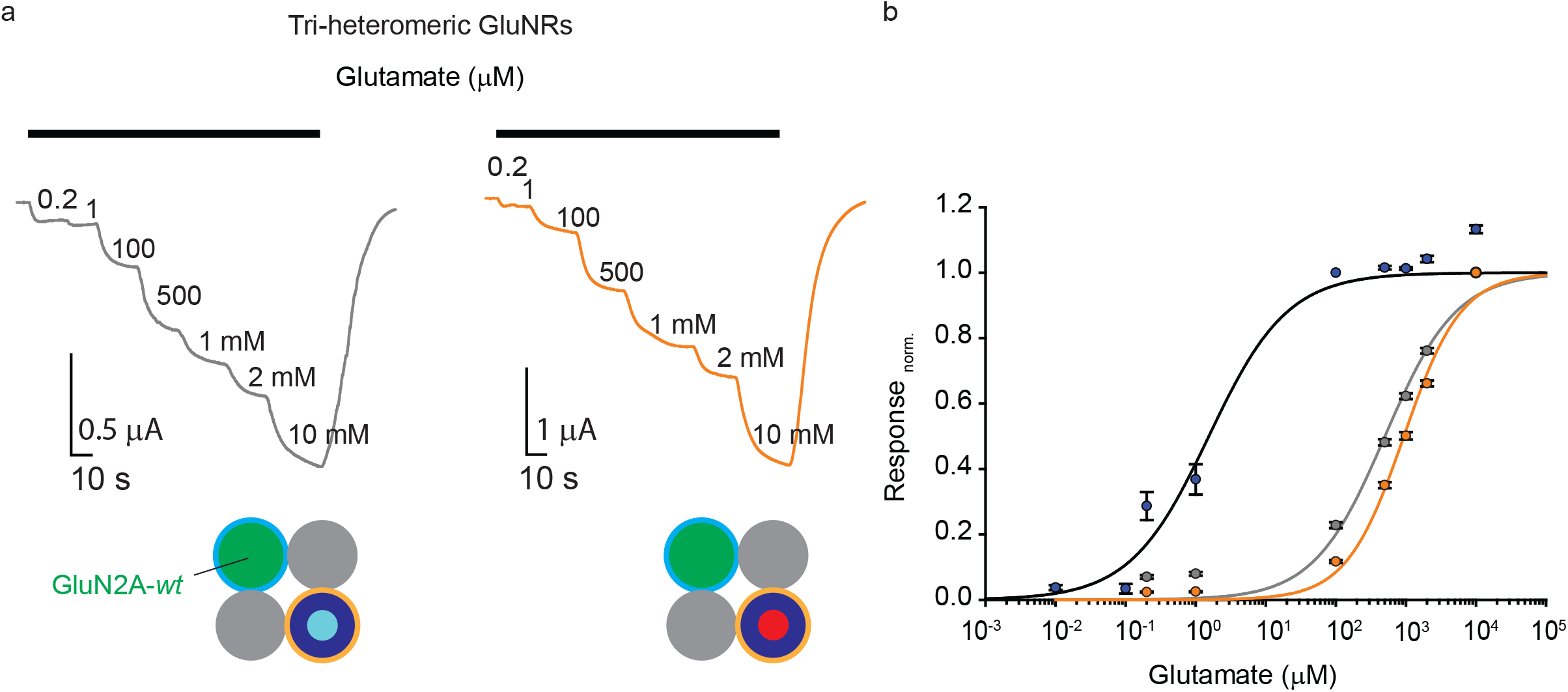
GluN2B-variants reduce glutamate potency of tri-heteromeric receptors. **a**. Representative traces from oocytes expressing tri–hetereomers composed of GluN1a-*wt* with GluN2A*wt*-C1 (green filled circle with cyan outline in bottom cartoon) and GluN2B-G689C-C2 (dark blue circle with cyan filled circle and orange outline in bottom cartoon) or with GluN2B-G689S-C2 (dark blue circle with red filled circle and orange outline) depicted in grey and orange respectively, in response to increasing glutamate concentrations. Glutamate (and glycine) application is noted by black bar above traces. Glutamate concentrations (in μM) are noted next to current steps. mM units are explicitly noted next to steps. **b**. Summary of dose-response curves for tri–hetereomeric GluN2B-receptors, colored-coded as in **(a)** and (**suppl 1.a)**.

### Mixed di- and tri-heteromeric channels containing the GluN2B-variants respond differently to GluN2B-selective potentiators

To assess whether channel activity could be rescued pharmacologically, we turned to examine spermine, a GluN2B-selective potentiator^33^. Of note, we were reluctant because purely di-heteromeric channels (containing two copies of the mutated subunits) do not respond to the drug^9^. However, the binding site for spermine lies at the dimer interface between the GluN1 and GluN2B subunits (at the N-terminal domains), and it remains debated whether a single intact interface is sufficient for its potentiation^33^. Remarkably, we found that *mixed* di-heteromeric receptors, composed of GluN2B*wt*-C1 assembled with GluN2B-G689C-C2 or GluN2B-G689S-C2, undergo robust potentiation by spermine, highly similar, although lower, to responses of *wt* receptors to the drug (**Fig. 4a, b**). Thus, GluN2B-G689C and GluN2B-G689S do not exert a dominant negative effect over spermine potentiation when assorted with a GluN2B-*wt* subunit. We also find that all GluN2A-*wt-*containing tri-heteromeric receptors were non-responsive to spermine (actually, slightly inhibited) under physiological pH and temperature (and -60 mV holding potential), regardless the identity of the GluN2B subunit (i.e., *wt* or mutated) (**Fig. 4a, b, Suppl. 1c**). These results show that potentiation by spermine indeed requires only a single intact interface between GluN1-*wt* and GluN2B-*wt* subunits (**Table 1**), however this does not hold true for tri-h eteromeric receptors. Thus, these results argue against the use of spermine as a possible treatment for GluN2B’s LoF mutations, at least postnatally, during which period tri-heteromeric receptors are the more abundant form. In fact, the use of spermine may actually worsen the clinical phenotype by further inhibiting extant tr-heteromers.

**Figure 4.**
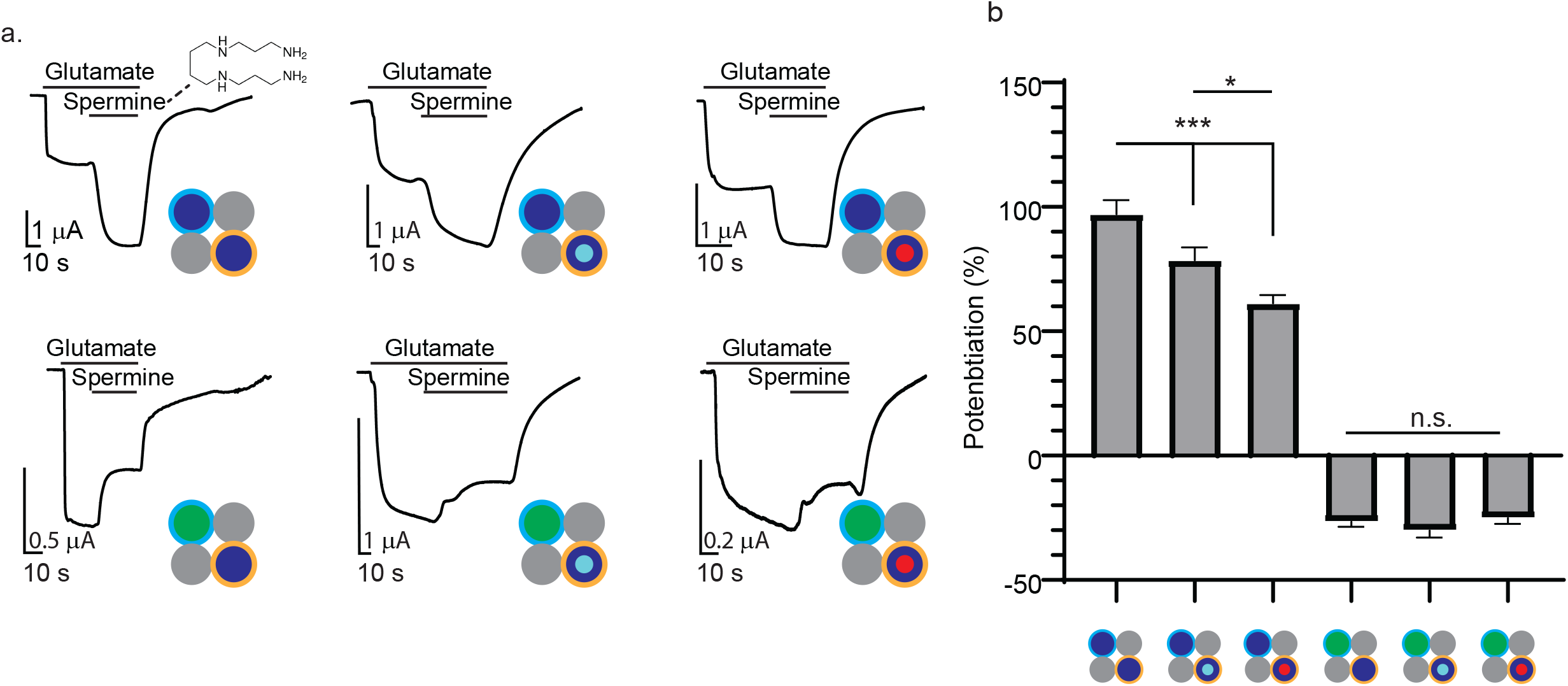
Spermine potentiates only *mixed* di-heteromeric receptors containing only one subunit of the GluN2B variants. **a**. Representative traces from oocytes expressing pure or *mixed* di-heteromers containing the GluN1a-*wt* subunit co-expressed with GluN2B*wt*-C1 and GluN2B*wt*-C2 or GluNB-G689C-C2 or G689S-C2 (top; left, middle and right traces, respectively) or tri-heteromers assembled from GluN1a-wt and GluN2A*wt-*C1 (green) with GluN2B*wt*-C2 (bottom left) or GluN2B-G689C-C2 (bottom middle) or GluN2B-G689S-C2 (bottom right) in response to 200 μM spermine (at pH 7.3) with 5 mM glutamate (+100 μM glycine) (black bars above traces). Molecular structure of spermine is shown. **b**. Summary of spermine potentiation (in %) of di- and tri-heteromers. *, p < 0.05; ***, p<0.001; n.s., not significant.

We proceeded to explore the effect of the neurosteroid pregnenolone-sulfate (PS), a well-established positive allosteric modulator (PAM) of di-heterometric GluN2A- or GluN2B-containing receptors^34,35^. Notably, and to the best of our knowledge, the use of PS on tri-heteromeric channels has yet to be explored. We found that purely *wt* or mixed di-heteromeric receptors containing GluN2B-subunits exhibit similar potentiation (∼200%) by 100 μM PS (**Fig. 5a, b, Suppl. 1d**). Unintuitively, all tri-heteromeric receptors (whether with a *wt* or a GluN2B variant) show similar potentiation (**Fig. 5b**) and, perhaps the most striking, purely mutated di-heteromeric receptors show the greatest potentiation (G689S-∼5-fold, G689C-∼4-fold, **Fig. 5b**). Thus, these clearly demonstrate that the binding domain of PS (within the transmembrane^35^) is not affected by the mutations in the LBD of the GluN2B subunits. Of note, the different extents of potentiation observed for the di-compared with tri-heteromers is suggested to stem from the diverging potentiation mechanisms of PS onto GluN2A and GluN2B subunits by PS^34^. Together, we show that PS is a powerful potentiator of purely mutated di-heteromers. We extend these findings towards tri-heteromers only to show, for the first time, that PS is a powerful potentiator of all channel types, regardless whether it includes a *wt* or mutated variant within the receptor complex. Thus, PS may serves to rescue the effects of the mutations by enhancing the currents of the receptors, as frequently suggested for di-heteromers^35–38^.

**Figure 5.**
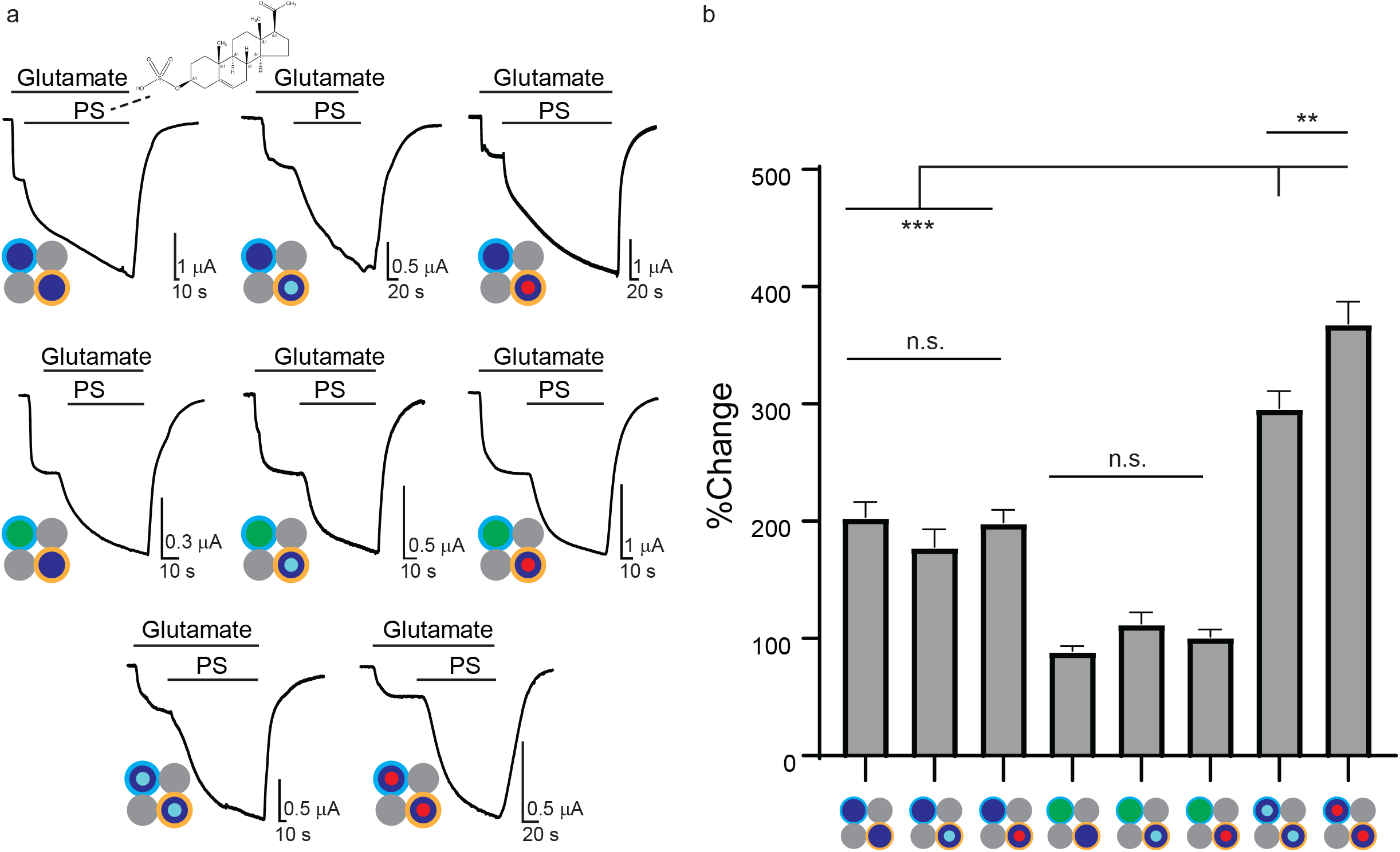
Pregnenolone-sulfate effectively potentiates all channel types regardless of presence of the GluN2B-variants. **a**. Representative traces from oocytes expressing pure or *mixed* di-heteromeric or tri-heteromeric GluNRs in response to 100 μM pregnenolone sulfate (PS) in the presence of 5 mM of glutamate (+100 μM glycine) (indicated by black bars above traces). Molecular structure of PS is shown in first panel. **B**. Summary of PS-potentiation (in %). **, p < 0.01; ***, p<0.001; n.s., not significant.

Lastly, we also examined the effect of Olanzapine, a derivative of clozapine, which has been previously suggested to act as a potential ‘enhancer’ of NMDARs^39–41^. Aside previous reports, additional motivation behind the examination of olanzapine is the fact that it is an FDA-approved anti-psychotic drug and therefore, if indeed active, would present novel opportunities to quickly obtain approval for treating *GRINopathies* at the clinic. Despite the latter, we found no evidence for any direct effect of three different physiologically-relevant concentrations of the drug over di- or tri-heteromeric receptors (**Suppl. 2a-e**), even though it did show its established effect (inhibition) over hERG channels (**Suppl. 2f**)^42^.

### PS rescues NMDAR-current amplitudes in cultured hippocampal neurons

We next turned to examine the functional outcome of the use of PS in neurons. We transfected cultured hippocampal neurons only with the different mutated GluN2B-subunits, relying on the endogenous subunits to assemble with, and traffic them to the membrane^43^. We first noted that overexpression of the variants caused a significant reduction in the NMDAR-current (**Fig. 6a, b- asterisk, and c**), as previously shown^9^, and— importantly—application of 100 μM PS increased the steady-state NMDA-dependent currents in neurons expressing the G689C and G689S variants to similar amplitudes to control neurons (**Fig.6b, c**). We also observed that potentiation by PS was most prominent in neurons overexpressing GluN2B-G689S (∼60%) (**Fig.6d**). Based on our oocytes recordings (see **Fig. 5**), this result may have suggested the over-abundance of purely di-heteromeric channels consisting of GluN2B-G689S variants at the membrane of neurons, although we ruled-this out by use of ifenprodil (a selective GluN2B-inhibitor^23,26^), showing that these neurons enclose the same fraction of GluN2B containing receptors (**Fig. 6e**).

**Figure 6.**
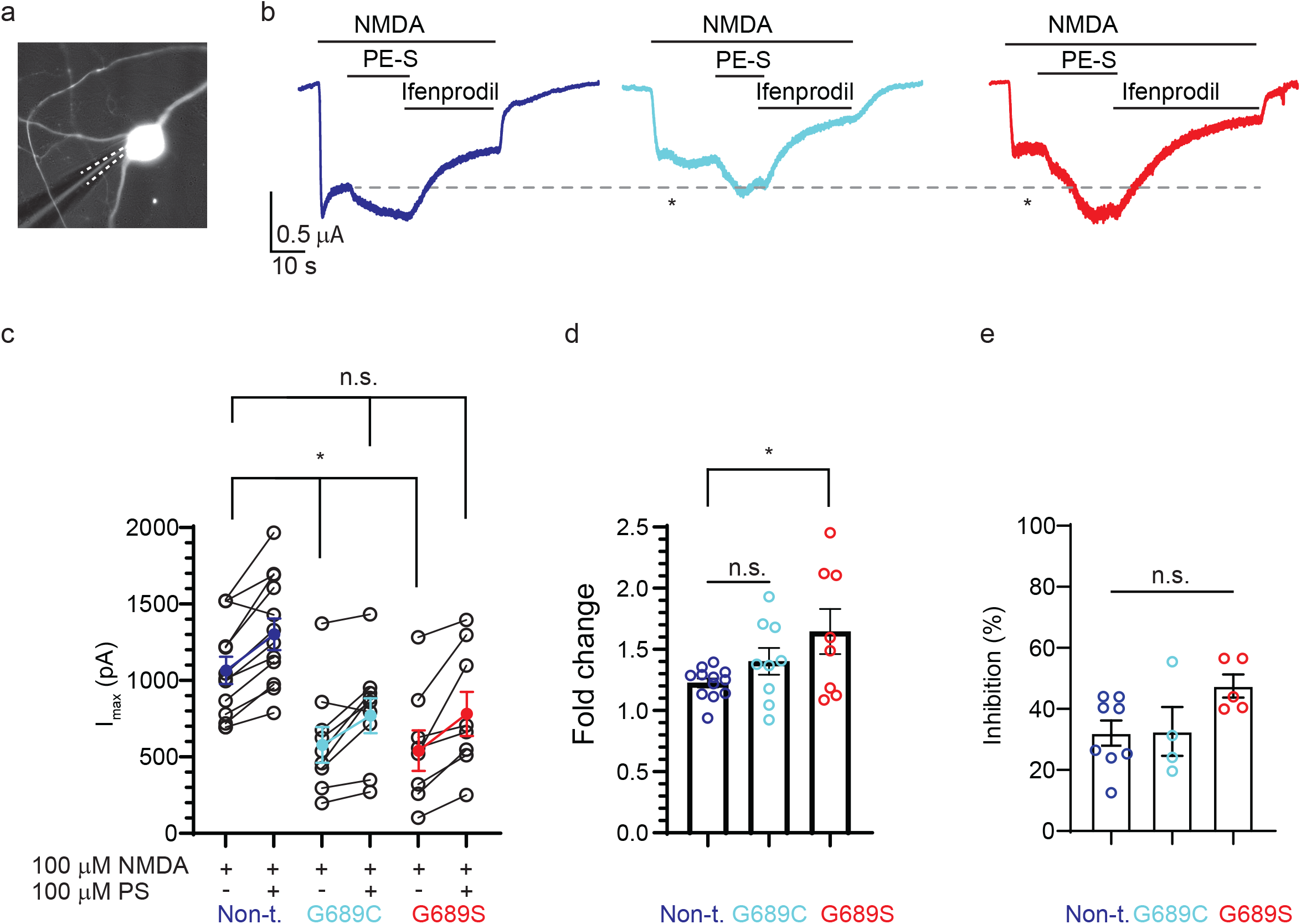
Pregnenolone-sulfate rescues NMDAR-current amplitudes in hippocampal neurons overexpressing GluN2B-variants. **a**. Representative micrograph of a hippocampal neuron overexpressing a GluN2B-variant and YFP (for visualization); recording pipette is highlighted by dashed white lines. **b**. Whole cell recordings of NMDAR-dependent currents from non-transfected (dark blue trace), GluN2B-G689C-transfected neuron (cyan) or GluN2B-G689S-transfected neurons (red) in response to 100 μM NMDA (+50 μM glycine) and 100 μM PS. The GluN2B-dependent fraction of the current was determined by application of 2.5 μM ifenprodil in the presence of the agonists. Note the strong and significant reduction in the maximal steady-state NMDAR-dependent currents between control and transfected neurons (dashed grey line and asterisks). **c**. Summary of the maximal currents (I_max_), before and after application of PS. Each bullet represents an individual neuron, with means (and SEM) depicted by blue (non-transfected), cyan (GluN2B-G689C) or red (GluN2B-G689S). **d**. Summary of the extant (fold change) of current potentiation by PS. **e**. Fraction of GluN2B-dependent current is unchanged between groups (% inhibition by ifenprodil). **, p < 0.01; n.s., not significant.

Collectively, our results demonstrate that the G689C- and G689S-variants reduce the total NMDAR-dependent currents, and instigate a very strong dominant negative effect over glutamate affinity of all forms of di- or tri-heteromeric receptors (**Figs. 1-3**). However, in the case of the use of spermine, only purely di-heteromeric channels containing the variants are non-responsive to the reagent, whereas *mixed* di-heteromers remain almost completely unperturbed (**Fig. 4**). The latter applies for PS as well, as all channel types are responsive to micromolar concentrations of the neurosteroid (**Figs. 5, 6**).

## Discussion

The tendency of NMDAR-subunits to assemble into tri-heteromeric receptors (i.e., two GluN1-subunits and two different GluN2 subunits, notably GluN2A and GluN2B) in the postnatal brain is widely gaining acceptance^3,21,44^. However, very little is known regarding the effect of single *GRIN* variants over the function of tri-heteromers, in particular at the synapse and their sensitivity (or irresponsiveness) to NMDAR-selective drugs (i.e.,^15,16,24,45,46^). This scenario limits our understanding of the *heterozygous*-nature of most *GRINopathies*, and is likely behind the very slow progression towards development of treatment to patients (e.g.,^12^).

Here, we make a small, albeit significant, step forward by examining the effect of two very extreme *de novo* analogous mutations in the context of di- and tri-heteromers, previously found in two pediatric patients (specifically, G689C and G689S^9^). We first focus on the most detrimental feature of both mutations over receptor function, namely their ultrapotent (∼2000-fold) reduction in glutamate affinity. We have previously shown that the mutated residue (G689) is precisely located at the opening of the LBD, which thereby affects the correct coordination and, consequently, binding of glutamate, yielding receptors with extremely high EC_50_ values (∼mM). Here, we show anew that a single GluN2B-variant, whether assembled with a GluN2B-*wt* subunit to form *mixed* di-heteromer or with a GluN2A-*wt*-subunit (i.e., tri-heteromer), strongly reduces the apparent glutamate affinity of the receptor, but the resulting affinities are less extreme than those obtained from purely di-heteromeric receptors containing two GluN2B-variants (**Figs. 1-3, and Table 1**). This was somewhat surprising, as NMDARs opening requires all four subunits to be *liganded* (i.e., occupied by a ligand) which implies that the least affine subunit should have dominated the final affinity of the receptor^29^. This can be explained by the well-established cooperativity observed between the subunits; each strengthening the affinity of the neighbors to their ligands^32^. Despite the latter, the negative effect of the two variants over mixed di- and tri-heteromeric receptors is strong enough so that their effectiveness is presumably below the expected glutamate concentrations that are readily obtained at the synaptic cleft, rendering both receptors and synapses oblivious (i.e., silent) to neurotransmission^9^. We would like to emphasize that this effect is not to be confused with cases of haploinsufficiency, for instance due to mutations that cause a drastic reduction in expression of synaptic receptors such as truncation mutations, even though they may share several functional similarities such as reduced currents at synspase^25,46,47^.

Another interesting, and counterintuitive finding is the effect of spermine, a GluN2B-selective potentiator. We have previously demonstrated that purely mutated di-heteromers are poorly responsive (or even inhibited in the case of the G689S variant) to spermine. However, when combined with a GluN2B-*wt* subunit, *mixed* di-heteromers regain their sensitivity to the reagent (**Fig. 4b**). This demonstrates that a single GluN2B-*wt* subunit is sufficient to reinstate proton and, consequently, spermine sensitivity, whereas glutamate affinity is governed by a single subunit (see above, **Figs. 1, 2**). Even more surprising, to which we and others have yet to find a plausible answer^21^, is the fact that *wt* tri-heteromers do not strongly respond to spermine despite the presence of an intact GluN2B-GluN1-interface. This likely stems from the dimer-of-dimer arrangement of the receptors, in which the transduction of the effect of spermine by the gating machinery requires coordinated action all dimers, and in tri-heteromers, there is one missing interface, which is not the case with the GluN2B variants. In fact, our findings further demonstrate that all tri-heteromers (whether with *wt* or variants) undergo slight, albeit significant, inhibition (-20%) by spermine (**Fig. 4b**). Our observations support the established consensus whereby spermine potentiation is absent in tri-heteromers (see review by Stroebel^7^), however the genuine inhibition observed here is unique and stands in contrast to previous reports (of which there are only two reports^26,44^). Closer scrutiny shows that the different effects seen heer may have resulted from very different experimental schemes, such as much higher acidity (pH =6.4), voltage (-30 mV) and lower glycine/spermine concentrations employed in previous reports; features which likely favor potentiation and/or mask inhibitions^48^. Regardless, these, show the challenge in trying to foresee how different variants may affect this highly complex and coordinated gating of NMDARs. Together, our results demonstrate that, despite the positive effect of spermine over *mixed* di-heteromers (which should be prevalent in prenatal stages), its use should be avoided postnatally as the largest population of receptors at synapses are likely tri-heteromers and these are readily inhibited by it (**Fig. 4b**).

We then proceeded to examine the suitability of a potentiator from the neurosteroids family of molecules, explicitly pregnenolone-sulfate (PS). Of note, and to the best of our knowledge, PS has never been assessed with tri-heteromers. We now show that PS effectively potentiates all receptor types, namely purely or *mixed* di-heteromeric receptors assembled with the various GluN2B-variants and all tri-heteromers tested (**Fig. 5**). In fact, purely di-heteromeric receptors (bearing two mutated GluN2B-subunits) exhibit significantly improved potentiation by PS compared to other receptor subtypes (**Fig. 5b**). These results are supported by our recordings of NMDAR-dependent currents in primary cultured neurons overexpressing the different GluN2B variants (**Fig. 6**), not to mention complement reports showing the ability of PS to enhance short- and long-term potentiation (STP^49,50^ and LTP^51–54^, respectively). Thus, and in spite of the observations above with spermine, the two GluN2B variants have no negative effect over the different receptors’ responses to PS. Yet, it should be noted that PS may suffer from lack of specificity, as it may jointly perform as a negative allosteric modulator (NAM) of GluN2C and GluN2D-containing di-heteromers^55^ (never tested on tri-heteromers) and of the GABA_A_-receptor^34^. The latter may promote susceptibility to epileptic seizures. Thus, owing to this the complex array of activities, whether PS could serve as a viable treatment for *GRINopathies* remains actively explored^32–34,54^. An alternative neurosteroid with similar potentiation capabilities towards GluN2B-receptorss is 24(S)-hydroxycholesterol (24-S), the major cholesterol metabolite in the brain^38^, to which a handful of synthetic analogues have been synthesized (e.g., SGE-201, SGE-301)^38^. Alternatively, there may be several means to increase the concentrations of 24(S) in the brain, for instance by promoting the activity of cytochrome P450 46A1 (CYP46A1) which converts cholesterol to 24-hydroxycholesterol, by FDA-approved drugs such as Efavirenz (an anti-retroviral compound)^56^. Despite this realization and potential benefit, we have yet to find a report that combines *GRINopathies* and Efavirenz, and we have yet to explore this strategy ourselves owing to lack of success in generating *GRIN2B*-G689C/S transgenic animals (see **Suppl. Text**).

In summary, we examined how a single variant affects the function of di- and tri-heteromers. We show that while a single dysfunctional subunit readily instigates a dominant negative effect over glutamate affinity, it does not dominate the effect over allosteric features of the receptor, notably potentiation by spermine and PS. Our results highlight the importance in examining how different mutations affect features in various receptor subtypes, as these cannot be deduced from observations from purely di-hetereomeric receptors. Lastly, and perhaps the most important, we show the feasibility of using a neurosteriod to rescue glutamatergic neurotransmission. Together, our study contributes to the ongoing efforts invested into the understanding of the pathophysiology of *GRINoparthies* variants, and provide insights into potential treatments.

## Methods

### *Xenopus laevis* oocytes extraction

*Xenopus laevis* oocytes were collected, processed, and injected with mRNA, as previously described^9,57^. Briefly, female frogs were anesthetized, and their ovaries were harvested. Ovaries were then treated with collagenase in ND96 Ca^+2^-free solution (in mM: 96 NaCl, 2 KCL, 1 MgCl2, 5 HEPES, pH=7.4) for 20 min at RT, to isolate and defolliculate the oocytes. Subsequently, oocytes were washed with ND96 Ca^+2^-free solution and stored in enriched ND96 medium (NDE) consisting of ND96 added with 2.5 mM sodium pyruvate, 1.8 mM CaCl_2_, 100 mg/ml streptomycin and 62.75 mg/ml penicillin. Lastly, hand-picked stage V oocytes were identified, separated, and incubated overnight at 18°, and then injected with mRNA.

### Dissociation, culturing, maintenance, and transfection of primary hippocampal neurons

Cultures of hippocampal primary neurons were established as previously stated^58^. Briefly, extracted rat neonates (P0) hippocampi were dissociated and plated onto 12 mm poly-D-lysine (Sigma-Aldrich, Cat. #P6407)-treated glass coverslips. Cultures were then maintained in an enriched growth media and grown at 37°C and 5% CO_2_. Following five days in-vitro (DIV), growth medium was supplemented with 4 μM cytosine-arabinoside (ARA-C) to suppress glia proliferation. At nine DIV, neurons were transfected using the calcium-phosphate method with 0.3 μg DNA of eYFP and 2 μg of GluN2B-G689C or GluN2B-G689S. Recordings were performed four to seven days past transfection.

### Molecular biology and *in vitro* mRNA preparation

Rat GluN1a-*wt*, rat GluN2A*wt*-C1/2 and rat GluN2B*wt*-C1/2 plasmids were obtained from Prof. Hansen K.B. (Montana University). Rat GluN2B-G689C-C1/2 and Rat GluN2B-G689S-C1/2 were generated using the QuikChange Site-Directed Mutagenesis Kit (Agilent,Cat. # 200518). Primers for GluN2B-G689C Mutagenesis: sense-5’-CGCTTTGGGACTGTGCCCAATTGCAGCACAGAGAGGAATATCCG -3’, antisense-5’-CGGATATTCCTCTCTGTGCTGCAATTGGGCACAGTCCCAAAGCG-3’; for GluN2B-G689S: sense-5’-CCGCTTTGGGACCGTGCCCAACAGCAGCACAGAGAGAAATATCCG-3’, antisense-5’-CGGATATTTCTCTCTGTGCTGCTGTTGGGCACGGTCCCAAAGCGG-3’. All PCR products were fully sequenced. For *in-vitro* mRNA transcription, plasmids were linearized with NotI, and transcription was obtained by mMessage-mMachine T7 kit (Thermo Scientific, Cat. #AM1344). Subsequently, mRNA concentrations were measured using a spectrophotometer. For selective expression of different compositions of GluNRs, mRNA of GluN1a was co-injected with GluN2A/B with different tails (C1 or C2) at a ratio of 1:3.75:3.75, at ∼28 ng mRNA/oocyte. For assessment of leak expression, mRNA of rat GluN1a was co-injected with GluN2Awt-C1 or GluN2Bwt-C1 (or both) at similar ratios and amounts as described above. Recordings were then performed 24-72 hrs after injection. For hERG expression, 25 ng mRNA/oocyte was injected.

### Two Electrode Voltage Clamp recordings in *Xenopus* laevis oocytes

Two electrode voltage clamp (TEVC) recordings were carried out 24–72 hrs after mRNA injections, as previously described^9,27^. We use a commercial amplifier (Warner Instruments, USA) and Digitizer (Digidata-1550B; Molecular Devices, USA), controlled by the pClamp10 software (Molecular Devices, USA). Electrode were made by pulling glass capillaries (Narishige, Japan) filled with 3 M KCl, into which we inserted chlorinated silver wires. Stage V oocytes were then clamped at -60 mV) and perfused with barth solution (in mM): 100 NaCl, 0.3, Bacl_2_, 5 HEPES, at pH=7.3 (adjusted by NaOH). Glutamate dose response experiments were performed with glutamate concentrations ranging between 0.2 μM and 10 mM (all containing 100 μM glycine). In the case of hERG channel recordings, oocytes were clamped at -60 mV for 1 s, followed by a voltage jump to 20 mV for 4 seconds, and an additional voltage jump to -50 mV for 6 seconds and return to baseline voltage, as previously described ^42^.

### Patch clamping of cultured neurons

We patched YFP-positive neurons at 13–16 DIV. YFP was excited by X-Cite LED illuminator (Excelitas Technologies). Electrode were made by pulling glass capillaries to resistance of 5-10 MΩ. Electrodes were filled with an intracellular solution containing (in mM): 135 K-gluconate, 10 NaCl, 10 HEPES, 2, MgCl_2_, 2 Mg^2+^-ATP, 1 EGTA, pH=7.3. Neurons were clamped at -70 mV. For assessing the total NMDAR-dependent current, neurons were perfused with extracellular solution containing (in mM): 138 NaCl, 10 Glucose, 5 HEPES, 2.5 CaCl_2_, 1.5 KCl, pH=7.4 (adjusted by NaOH), and (μM): 50 glycine and 20 CNQX (Alomone labs, Cat #: C-140), 10 gabazine (Alomone labs, Cat #: G-215), 1 TTX (Alomone labs, Cat #: T-550), then total amplitude was obtained by application of 100 μM of NMDA (Alomone labs, Cat #: N-170) for exclusive activation of GluNRs. Potentiation was assessed by perfusing cells with the extracellular solution along with 100 μM NMDA and pregnenolone sulfate (Cat. #P162). Lastly, Inhibition of GluN2B-containing receptors was evaluated by application of 2.5 μM ifenprodil (Alomone labs, Cat #: I-105)

### Potentiation of NMDAR-currents by drugs

Spermine (Cat. #S3256), Pregnenolone-sulfate (Cat. #P162) and olanzapine (Cat. #O1141) were purchased from Sigma-Aldrich. 200 mM spermine stock solution (in water) was freshly made on the day of the experiment. 50 mM and 10 mM stock solutions of Pregnenolone-sulfate and olanzapine, respectively, were made in DMSO. Potentiation was evaluated by application of the drugs in the presence of 5 mM glutamate and 100 μM glycine. When assessing potentiation with Pregnenolone-sulfate, all other solutions also included equal amounts of the DMSO carrier.

### Data and Statistical analysis

Electrophysiology data were analyzed by Clampfit (Molecular Devices, USA), plotted using GraphPad 8 or SigmaPlot 11. EC_50_ values were extracted by fitting the data to Hill equation: Response = 1/ (1+ [(glutamate)/EC_50_]^nH^). All data are shown as mean ± SEM. In all experiment N indicates the number of independent experiments, whereas n indicates the number of cells recorded. Statistical significance was assessed by using one-way ANOVA for multiple group comparison and *post hoc* Tukey test. n.s., non-significant; *, p<0.05; **, p< 0.01; ***, p<0.001.

### Ethics

The Technion Institutional Animal Care and Use Committee approved all experiments with *Xenopus laevis* oocytes (permit SB, no. IL-162-10-25), rat hippocampal neurons (permit SB, no. IL-143-12-20) and for the design of a transgenic mouse model bearing the GRIN2B-G689C patient-derived mutation (permit SB, no. IL-125-11-20).

## Acknowledgements

We thank Dr Hansen Kasper B. for providing us the plasmids for selective expression of GluNRs, and Alomone labs (Jerusalem, Israel) for their kind support with providing us reagents and compounds. The research submitted is in partial fulfillment for a doctoral degree for SK

## Legends

**Supplementary Figure 1. Assessing leak expression for tri-heteromers**.

**a**. Cartoon depiction of a surface expression-enabled tri-heteromeric receptor composed of two wildtype GluN1a subunits (gray) assembled with one GluN2A*wt*-C1 subunit (green with cyan outline) and one GluN2B*wt*-C2 subunit (dark blue with orange outline) and a representative trace (middle trace). Right panel shows a representative trace of leak tri-heteromeric current recorded from oocytes co-expressing GluN1a-wt+GluN2A*wt*-C1+GluN2B-*wt*-C1. Glutamate (and glycine) application is noted by black bar above traces. **b**. Summary of normalized currents (I_max_) from glutamate dose response of tri-heteromeric receptors recorded in two independent experiments 72 hr. after injection. Median is highlighted in red . **c**. Summary of normalized currents (I_max_) from potentiation experiments recorded in two independent experiments 24 hr. after injection, with the median highlighted in red. ***, p < 0.001;

**Supplementary Figure 2. Olanzapine does not directly potentiate GluNRs**.

**a-e**. Representative traces from oocytes expressing di- and tri-heteromers and their responses to three different concentrations of olanzapine (indicated next to current steps on trace) and summary (right). Glutamate (and glycine) application as well as olanzapine is indicated by black bars above traces. **f**. Representative trace from an oocyte expressing the hERG channel, before (black) and after (red) application of 20 μM olanzapine and summary of inhibition (right).

## References

1. Hardingham, G. E. & Bading, H. Synaptic versus extrasynaptic NMDA receptor signalling: implications for neurodegenerative disorders. Nat. Rev. Neurosci. 11, 682–696 (2010).

2. Paoletti, P., Bellone, C. & Zhou, Q. NMDA receptor subunit diversity: impact on receptor properties, synaptic plasticity and disease. Nat. Rev. Neurosci. 14, 383–400 (2013).

3. Hansen, K. B. et al. Structure, Function, and Pharmacology of Glutamate Receptor Ion Channels. Pharmacol. Rev. 73, 1469–1658 (2021).

4. Gonda, S. et al. GluN2B but Not GluN2A for Basal Dendritic Growth of Cortical Pyramidal Neurons. Front. Neuroanat. 14, (2020).

5. Zhang, Z., Peterson, M. & Liu, H. Essential role of postsynaptic NMDA receptors in developmental refinement of excitatory synapses. Proc. Natl. Acad. Sci. 110, 1095–1100 (2013).

6. Kelsch, W., Li, Z., Eliava, M., Goengrich, C. & Monyer, H. GluN2B-Containing NMDA Receptors Promote Wiring of Adult-Born Neurons into Olfactory Bulb Circuits. J. Neurosci. 32, 12603–12611 (2012).

7. Stroebel, D., Casado, M. & Paoletti, P. Triheteromeric NMDA receptors: from structure to synaptic physiology. Curr. Opin. Physiol. 2, 1–12 (2018).

8. Endele, S. et al. Mutations in GRIN2A and GRIN2B encoding regulatory subunits of NMDA receptors cause variable neurodevelopmental phenotypes. Nat. Genet. 42, 1021–1026 (2010).

9. Kellner, S. et al. Two de novo GluN2B mutations affect multiple NMDAR-functions and instigate severe pediatric encephalopathy. eLife 10, e67555 (2021).

10. Addis, L. et al. Epilepsy-associated GRIN2A mutations reduce NMDA receptor trafficking and agonist potency – molecular profiling and functional rescue. Sci. Rep. 7, 66 (2017).

11. Amador, A. et al. Modelling and treating GRIN2A developmental and epileptic encephalopathy in mice. Brain (2020) doi:10.1093/brain/awaa147.

12. Soto, D. et al. L-Serine dietary supplementation is associated with clinical improvement of loss-of-function GRIN2B-related pediatric encephalopathy. Sci. Signal. 12, (2019).

13. García-Recio, A. et al. GRIN database: A unified and manually curated repertoire of GRIN variants. Hum. Mutat. 42, 8–18 (2021).

14. Platzer, K. et al. GRIN2B encephalopathy: novel findings on phenotype, variant clustering, functional consequences and treatment aspects. J. Med. Genet. 54, 460–470 (2017).

15. Swanger, S. A. et al. Mechanistic Insight into NMDA Receptor Dysregulation by Rare Variants in the GluN2A and GluN2B Agonist Binding Domains. Am. J. Hum. Genet. 99, 1261–1280 (2016).

16. Li, J. et al. De novo GRIN variants in NMDA receptor M2 channel pore-forming loop are associated with neurological diseases. Hum. Mutat. 40, 2393–2413 (2019).

17. Xu, X.-X. & Luo, J.-H. Mutations of N-Methyl-D-Aspartate Receptor Subunits in Epilepsy. Neurosci. Bull. 34, 549–565 (2018).

18. XiangWei, W., Jiang, Y. & Yuan, H. De novo mutations and rare variants occurring in NMDA receptors. Curr. Opin. Physiol. 2, 27–35 (2018).

19. Myers, S. J. et al. Distinct roles of GRIN2A and GRIN2B variants in neurological conditions. Preprint at https://doi.org/10.12688/f1000research.18949.1 (2019).

20. Vieira, M., Yong, X. L. H., Roche, K. W. & Anggono, V. Regulation of NMDA glutamate receptor functions by the GluN2 subunits. J. Neurochem. 154, 121–143 (2020).

21. Stroebel, D., Casado, M. & Paoletti, P. Triheteromeric NMDA receptors: from structure to synaptic physiology. Curr. Opin. Physiol. 2, 1–12 (2018).

22. Tovar, K. R., McGinley, M. J. & Westbrook, G. L. Triheteromeric NMDA Receptors at Hippocampal Synapses. J. Neurosci. 33, 9150–9160 (2013).

23. Hansen, K. B., Ogden, K. K., Yuan, H. & Traynelis, S. F. Distinct Functional and Pharmacological Properties of Triheteromeric GluN1/GluN2A/GluN2B NMDA Receptors. Neuron 81, 1084–1096 (2014).

24. Han, W. et al. Opportunities for Precision Treatment of GRIN2A and GRIN2B Gain-of-Function Variants in Triheteromeric N-Methyl-D-Aspartate Receptors. J. Pharmacol. Exp. Ther. 381, 54–66 (2022).

25. Elmasri, M. et al. Synaptic Dysfunction by Mutations in GRIN2B: Influence of Triheteromeric NMDA Receptors on Gain-of-Function and Loss-of-Function Mutant Classification. Brain Sci. 12, 789 (2022).

26. Stroebel, D., Carvalho, S., Grand, T., Zhu, S. & Paoletti, P. Controlling NMDA Receptor Subunit Composition Using Ectopic Retention Signals. J. Neurosci. 34, 16630–16636 (2014).

27. Berlin, S. et al. Two Distinct Aspects of Coupling between Gαi Protein and G Protein-activated K+ Channel (GIRK) Revealed by Fluorescently Labeled Gαi3 Protein Subunits. J. Biol. Chem. 286, 33223–33235 (2011).

28. Neame, S. et al. The NMDA receptor activation by d-serine and glycine is controlled by an astrocytic Phgdh-dependent serine shuttle. Proc. Natl. Acad. Sci. 116, 20736–20742 (2019).

29. Kussius, C. L. & Popescu, G. K. Kinetic basis of partial agonism at NMDA receptors. Nat. Neurosci. 12, 1114–1120 (2009).

30. Johnson, J. W. & Ascher, P. Glycine potentiates the NMDA response in cultured mouse brain neurons. Nature 325, 529–531 (1987).

31. Kleckner, N. W. & Dingledine, R. Requirement for glycine in activation of NMDA-receptors expressed in Xenopus oocytes. Science 241, 835–837 (1988).

32. Sun, W., Hansen, K. B. & Jahr, C. E. Allosteric interactions between NMDA receptor subunits shape the developmental shift in channel properties. Neuron 94, 58-64.e3 (2017).

33. Mony, L., Zhu, S., Carvalho, S. & Paoletti, P. Molecular basis of positive allosteric modulation of GluN2B NMDA receptors by polyamines. EMBO J. 30, 3134–3146 (2011).

34. Geoffroy, C., Paoletti, P. & Mony, L. Positive allosteric modulation of NMDA receptors: mechanisms, physiological impact and therapeutic potential. J. Physiol. 600, 233–259 (2022).

35. Hrcka Krausova, B. et al. Site of Action of Brain Neurosteroid Pregnenolone Sulfate at the N-Methyl-D-Aspartate Receptor. J. Neurosci. 40, 5922–5936 (2020).

36. Kysilov, B. et al. Pregnane-based steroids are novel positive NMDA receptor modulators that may compensate for the effect of loss-of-function disease-associated GRIN mutations. Br. J. Pharmacol. 179, 3970–3990 (2022).

37. Vyklicky, V. et al. Surface Expression, Function, and Pharmacology of Disease-Associated Mutations in the Membrane Domain of the Human GluN2B Subunit. Front. Mol. Neurosci. 11, 110 (2018).

38. Paul, S. M. et al. The Major Brain Cholesterol Metabolite 24(S)-Hydroxycholesterol Is a Potent Allosteric Modulator of N-Methyl-D-Aspartate Receptors. J. Neurosci. 33, 17290–17300 (2013).

39. Arvanov, V. L., Liang, X., Schwartz, J., Grossman, S. & Wang, R. Y. Clozapine and haloperidol modulate N-methyl-D-aspartate- and non-N-methyl-D-aspartate receptor-mediated neurotransmission in rat prefrontal cortical neurons in vitro. J. Pharmacol. Exp. Ther. 283, 226–234 (1997).

40. Tanahashi, S., Yamamura, S., Nakagawa, M., Motomura, E. & Okada, M. Clozapine, but not haloperidol, enhances glial D-serine and L-glutamate release in rat frontal cortex and primary cultured astrocytes. Br. J. Pharmacol. 165, 1543–1555 (2012).

41. Wittmann, M., Marino, M. J., Henze, D. A., Seabrook, G. R. & Conn, P. J. Clozapine Potentiation of N-Methyl-d-aspartate Receptor Currents in the Nucleus Accumbens: Role of NR2B and Protein Kinase A/Src Kinases. J. Pharmacol. Exp. Ther. 313, 594–603 (2005).

42. Lee, H. J., Choi, J.-S. & Hahn, S. J. Mechanism of inhibition by olanzapine of cloned hERG potassium channels. Neurosci. Lett. 609, 97–102 (2015).

43. Berlin, S. et al. A family of photoswitchable NMDA receptors. eLife 5, e12040 (2016).

44. Cheriyan, J., Balsara, R. D., Hansen, K. B. & Castellino, F. J. Pharmacology of triheteromeric N-Methyl-d-Aspartate Receptors. Neurosci. Lett. 617, 240–246 (2016).

45. Amin, J. B., Leng, X., Gochman, A., Zhou, H.-X. & Wollmuth, L. P. A conserved glycine harboring disease-associated mutations permits NMDA receptor slow deactivation and high Ca2+ permeability. Nat. Commun. 9, 3748 (2018).

46. Santos-Gómez, A. et al. Disease-associated GRIN protein truncating variants trigger NMDA receptor loss-of-function. Hum. Mol. Genet. 29, 3859–3871 (2021).

47. Liu, S. et al. A Rare Variant Identified Within the GluN2B C-Terminus in a Patient with Autism Affects NMDA Receptor Surface Expression and Spine Density. J. Neurosci. 37, 4093–4102 (2017).

48. Traynelis, S. F. et al. Glutamate Receptor Ion Channels: Structure, Regulation, and Function. Pharmacol. Rev. 62, 405–496 (2010).

49. Schiess, A. R. & Partridge, L. D. Pregnenolone sulfate acts through a G-protein-coupled sigma1-like receptor to enhance short term facilitation in adult hippocampal neurons. Eur. J. Pharmacol. 518, 22–29 (2005).

50. Schiess, A. R. B., Scullin, C. S. & Partridge, L. D. Neurosteroid-induced enhancement of short-term facilitation involves a component downstream from presynaptic calcium in hippocampal slices. J. Physiol. 576, 833–847 (2006).

51. Sliwinski, A., Monnet, F. p., Schumacher, M. & Morin-Surun, M. p. Pregnenolone sulfate enhances long-term potentiation in CA1 in rat hippocampus slices through the modulation of N-methyl-D-aspartate receptors. J. Neurosci. Res. 78, 691–701 (2004).

52. Sabeti, J., Nelson, T. E., Purdy, R. H. & Gruol, D. L. Steroid pregnenolone sulfate enhances NMDA-receptor-independent long-term potentiation at hippocampal CA1 synapses: Role for L-type calcium channels and sigma-receptors. Hippocampus 17, 349–369 (2007).

53. Chen, L., Miyamoto, Y., Furuya, K., Mori, N. & Sokabe, M. PREGS induces LTP in the hippocampal dentate gyrus of adult rats via the tyrosine phosphorylation of NR2B coupled to ERK/CREB [corrected] signaling. J. Neurophysiol. 98, 1538–1548 (2007).

54. Chen, L. et al. Modulatory metaplasticity induced by pregnenolone sulfate in the rat hippocampus: A leftward shift in LTP/LTD-frequency curve. Hippocampus 20, 499–512 (2010).

55. Malayev, A., Gibbs, T. T. & Farb, D. H. Inhibition of the NMDA response by pregnenolone sulphate reveals subtype selective modulation of NMDA receptors by sulphated steroids. Br. J. Pharmacol. 135, 901–909 (2002).

56. Petrov, A. M. et al. CYP46A1 Activation by Efavirenz Leads to Behavioral Improvement without Significant Changes in Amyloid Plaque Load in the Brain of 5XFAD Mice. Neurotherapeutics 16, 710–724 (2019).

57. Berlin, S. et al. A Collision Coupling Model Governs the Activation of Neuronal GIRK1/2 Channels by Muscarinic-2 Receptors. Front. Pharmacol. 11, (2020).

58. Berlin, S. & Isacoff, E. Y. Optical Control of Glutamate Receptors of the NMDA-Kind in Mammalian Neurons, with the Use of Photoswitchable Ligands. in Biochemical Approaches for Glutamatergic Neurotransmission (eds. Parrot, S. & Denoroy, L.) vol. 130 293–325 (Springer New York, 2018).

